# Estrogen Signaling During Abrupt Involution Leads to Long-Term Metabolic Dysfunction Similar to Estrogen Receptor Negative Breast Cancer

**DOI:** 10.1101/2025.10.09.681352

**Authors:** Kate Ormiston, Neelam Shinde, Gautam Sarathy, Allen Zhang, Morgan Bauer, Rajni Kant Shukla, Sara Alsammerai, Annapurna Gupta, Djawed Bennouna, Julia Wesolowski, Xiaoli Zhang, Rachel E. Kopec, Eswar Shankar, Kristin I. Stanford, Ramesh K. Ganju, Sarmila Majumder, Bhuvaneswari Ramaswamy, Daniel G. Stover

## Abstract

Epidemiological data links lack of breastfeeding with increased risk of breast cancer. Breast tissue undergoes remodeling to pre-pregnancy state after birth through involution. Long-term breastfeeding leads to gradual involution (GI). Lack of breastfeeding leads to abrupt involution (AI). While estrogen impacts repopulation of adipocytes, AI causes several precancerous changes in the mouse mammary gland. The impact of AI on adipocyte repopulation and metabolism is yet to be elucidated.

**Objectives:** To investigate effects of AI on mammary gland metabolism and its potential link to breast cancer.

**Methods:** At partum (day 0), FVB/n dams were randomized to AI or GI and standardized to 6 pups. AI mice had pups removed on day 7 postpartum to mimic short-term breastfeeding. GI mice had 3 pups each were removed on day 28 and 31 postpartum to mimic gradual weaning. Mammary glands were harvested on day 28, 56, and 120 postpartum. Subset of AI mice had long-term sustained release tamoxifen placed subscapular on day 8 postpartum. Metabolic changes were assessed using: 1) transcriptional; 2) functional; 3) oxidative stress; and 4) metabolites analysis.

**Results:** Day 28 GI glands sustains/continues milk synthesis pathways impacting metabolic comparison with day28 AI glands. Day 28 AI when compared to day 56 GI showed upregulation of estrogen signaling, neutrophil degranulation, glucose metabolism, RNA synthesis, and down regulation of adipogenesis and glycolysis. At day 120, AI glands had downregulation of oxidative phosphorylation and upregulation of mitochondria dysfunction similar to pregnancy associated estrogen receptor negative breast cancer. Tamoxifen treatment of AI dams showed metabolic pathways and estrogen signaling similar to that of GI glands on day 28.

**Conclusion:** Early metabolic phenotypes in AI and GI glands may be caused by differences in adipocyte repopulation related to estrogen signaling. Long-term metabolic effects of AI lead to similar metabolic effects found in breast cancer.

## 1. Introduction

Breast cancer is the most prevalent cancer among women in the United States, with 1 in 8 women being diagnosed [1]. Epidemiological studies have linked lack of breastfeeding with increased risk of breast cancer, specifically triple negative breast cancer (TNBC) [2–6]. Large collaborative analysis from multiple epidemiological studies comprising over 50,000 breast cancer cases and controls concluded for 12 months of breastfeeding, breast cancer risk was reduced by 4.3% [5]. Several studies show 25-50% lower risk of TNBC in women who breastfed four to six months compared to parous women who never breastfed [4]

The mammary gland has remarkable ability to rapidly proliferate and shift to lactation during pregnancy [7–10]. At completion of breastfeeding, the mammary gland remodels to near pre-pregnancy state through involution [7]. Long-term breastfeeding and gradual weaning leads to a gradual remodeling process; gradual involution (GI) [10]. Abrupt discontinuation of breastfeeding or lack of initiation after birth leads to abrupt involution (AI) [10]. While the epidemiological link between lack of breastfeeding and breast cancer is clear [2–5], the mechanisms remain poorly understood. Research examining lack of breastfeeding following full-term pregnancy has suggested involution as the critical window involved in risk of breast cancer [7–10].

Mouse models of AI resulted in a pro-tumorigenic environment in the mammary gland [8–10]. Studies show short term effects of AI including greater cell death and turnover, heightened immune response, and tissue remodeling likened to wound healing [7–11]. AI leads to increased and sustained inflammation over prolonged periods, increased collagen deposition, and estrogenic signaling [10]. The mammary gland of mice that underwent AI displayed long-term increased cell proliferation, hyperplasia, and squamous metaplasia [10]. These studies conclude that AI leads to pre-cancerous changes within the mammary gland [7–11].

During AI estradiol binds to estrogen receptor alpha (ERα) causing an increase in neutrophils and extracellular remodeling which cause adipocyte re-population during the second phase of involution [12]. Adipocytes in AI glands expand after differentiation due to hypertrophy of excess milk lipids [9]. Research has shown that rapid adipocyte re-population and hypertrophy cause hypoxia and metabolic reprogramming [13]. Breastfeeding and breast cancer are two highly metabolic processes. However, the changes in metabolism related to AI and GI within the mammary gland have not been studied. We hypothesized that metabolic alterations induced by AI versus GI may drive resultant phenotypic differences in hyperplasia and inflammation. The objective of this study was to evaluate the short-term and long-term impact of AI on mammary gland metabolism using transcriptomic and functional analyses.

Our study shows similar AI adipocyte repopulation described in previous studies but after completion of involution and the type of involution causes differences in adipocyte repopulation. Accompanied with adipocyte repopulation AI and GI processes lead to differences in metabolic outcomes with long-term AI mammary glands mimicking metabolic phenotype of estrogen receptor negative (ER-) pregnancy associated breast cancer (PABC).

## 2. Methods

### Animals

All experiments were conducted according to National Institute of Health Guide for the Care and Use of Laboratory Animals. All protocols were approved by the Ohio State Institutional Animal Care and Use Committee. FVB/n female mice from Jackson Laboratory (Bar Harbor, ME) were paired for breeding at 8 weeks old. Female mice were separated prior to giving birth and housed individually. On day 0 (birth) mice were standardized to 6 pups per dam. On day 7 postpartum, mice were randomized to AI or GI. For AI, pups were removed from dam on day 7. For GI, 3 pups were removed from dam on day 28 and the remaining 3 pups were removed on day 31. Our model utilizes an extended weaning procedure for GI mice (day 28-31) in an effort to mimic long-term breastfeeding in accordance with the World Health Organizations (WHO) recommendations of breastfeeding for two years [14]. Research supports that rodent offspring will continue to periodically nurse when housed with the mother, even though offspring predominantly consume solid food [15,16]. Mice were harvested on 28-, 56-, or 120-days post-partum (**Figure 1a**).

**Figure 1.**
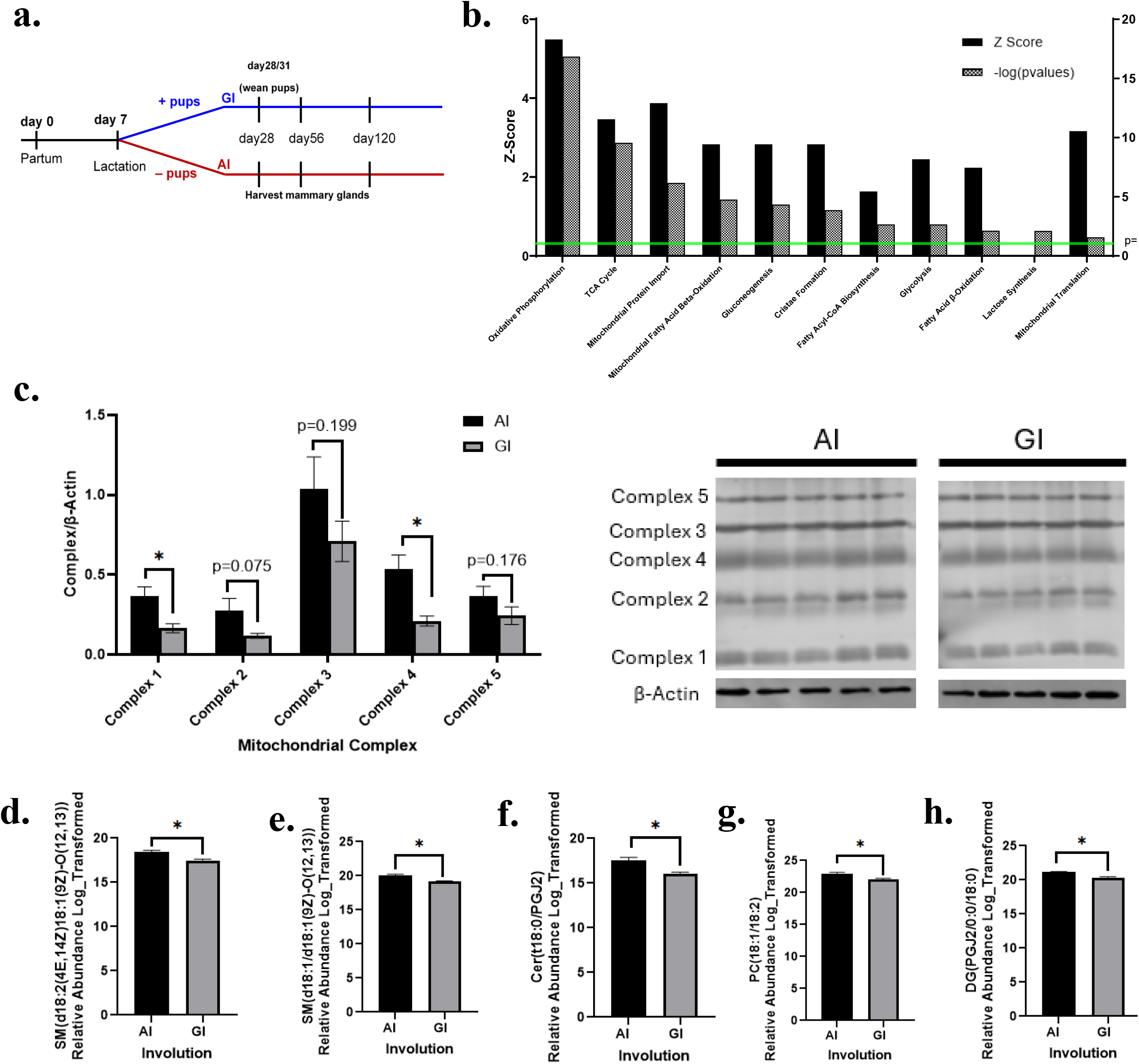
Metabolic Effects of Abrupt vs Gradual Involution 28 Days Post-partum. **a**. Experimental timeline. **b**. Ingenuity pathway analysis of metabolic pathways significantly impacted by AI. **c**. Western blot analysis of mitochondrial complexes. **d-h**. Significantly increased lipid species in AI glands. AI= abrupt involution. GI=gradual involution. Data presented as mean ± SEM. *depicts significant differences (p<0.05) between AI and GI glands.

### Treatment with Tamoxifen

A subset of FVB/n mice underwent breeding and AI. On day 8 postpartum, AI mice had either sustained release tamoxifen citrate (5mg) or placebo pellet placed in the subscapular region for 21 days. Mice were harvested on day 28.

### RNA Extraction

Mammary gland (50 mg piece) was cut and homogenized in Trizol using Precellys Lysing Kit (Bertin Technologies, France) and Precellys Evolution Homogenizer. Total RNA was isolated following Trizol-RNA isolation protocol. RNA was subjected to clean-up and concentration (Norgen Kit #23600; Biotek Corp., Canada) per protocol. RNA concentration and purity were determined using Nanodrop (ThermoFisher, Waltham, MA).

### Gene Expression Analysis

RNA extracted from mammary glands on days 28, 56, and 120 were analyzed via Affymetrix Clairom D Mouse Assay (Applied Biosystem; Waltham, MA) by Genomic Shared Resource at Ohio State University Comprehensive Cancer Center. Quality and purity were validated through tapestation analysis (n=3/group). Data was quality checked, and gene expression was analyzed using Transcription Analysis Console software (ThermoFisher).

### Ingenuity Pathway Analysis

Ingenuity Pathway Analysis software (Qiagen, Hilden, Germany) was used to compare gene expression data between AI and GI for impacted pathways. Pathways were selected based on two criteria: metabolism and energy production. Pathways having a p-value <0.05 were reported.

### qPCR

Total RNA extracted from mammary glands on days 28, 56, and 120 was utilized to synthesize cDNA using High Capacity cDNA Reverse Transcription kit (Applied Biosystems). qPCR was run (n=4-5 per group) using SYBR green (Biorad; Hercules, CA). Primers were purchased through Millipore-Sigma (Burlington, MA). Primer sequences provided in **Supplementary Table 1**. Primer fidelity was confirmed using agarose gel electrophoresis.

### Western Blot

Mammary gland samples (n=4-5; 50mg) added to RIPA buffer with 0.1M PMSF, 1% v/v phosphatase inhibitor cocktail 2 (P-5726; Sigma-Aldrich, St. Louis, MO), phosphatase inhibitor cocktail 3 (P0044-5; Sigma-Aldrich), and protease inhibitor (P8340; Sigma-Aldrich). Tissues were homogenized using Precellys lysing, allowed to sit on ice for 30 minutes, and centrifuged at 16100 x G for 10 minutes at 4 degrees Celsius. Protein was estimated by Pierce BCA protein assay kit #23225 (ThermoFisher). The mammary gland lysates were separated by SDS-PAGE buffer and transferred to PVDF membranes. Protein was immunoblotted with primary anti-bodies: Ox-phos Rodent WB Cocktail (45-8099; Invitrogen; Waltham, MA). Blots were incubated with corresponding secondary antibodies IRDye 680RD donkey anti-mouse IgG or IRDye 800CW goat anti-rabbit IgG and developed Odyssey CLX (Licor; Lincoln, Nebraska). Images were quantified via Image Studio Version 5.3 (Licor). Full blots are available in **Supplementary Figure 1**.

### Immunohistochemistry (IHC)

Mammary glands (n=3-5/group) were fixed in 10% neutral-buffered formalin for 72 hours and paraffin embedded. Ten micron sections were cut and fixed on glass slides. Sections were stained utilizing anti-F4/80 (1:500 Invitrogen MF48000) and anti-ERα (1:2000, Abcam ab32063). For adipocytes measurements, F4/80 slides were imaged used EVOS 5700 microscope (Thermo Fisher Scientific, Waltham, MA) and adipocytes quantified using ImageJ (NIH). ERα slides were analzyed using Vectra Microscope (Akoya Biosciences, Marlborough, MA) and InForm Software (Akoya Biosciences).

### Targeted Metabolomics

Mammary glands (50mg) were processed for metabolomics analyses using Liquid Chromatography-column isolation. Samples were weighed, and extraction solution added (MeOH:Chloroform 1:1) at 200 mg/ml. Extractions were performed using C18 column and Hilic column to capture non-polar and polar metabolites. Metabolomics measurements were performed using QTOF. Analyses were performed using XCMS online tools. Labelled internal standards for lactic acid, succinic acid, citric acid and palmitic acid (Cambridge Isotope Laboratories, Inc) were used to quantify metabolites in the mammary gland (n=4-5/group). Samples were aligned with a score of 90% or above and upon feature detection, ANOVA p-values between groups were calculated with a cutoff of 0.05. With database matching using the Human Metabolome Database, selecting for adducts M+H, M+Na, M+K, M+2H, for positive mode or M-H, M+Cl, and M-2H in negative mode and less than 10 ppm mass error, unique features were tentatively identified as potential metabolites.

### Sample Extraction for Untargeted Lipidomics

The extraction protocol was adapted from a previously described method [13]. Lipids were extracted from mammary gland with n=5 per group. Mammary gland (25 mg) was mixed with Optima methanol (250 µL), type I deionized water (125 µL) and zircona beads (0.5 mm). Samples were homogenized using a Mini-bead beater-16 (Biospec Products, Tulsa, OK) for 30 seconds, then vortexed for 1 min. Afterwards, 1000 µL of HPLC-grade methyl-tert butyl ether (MTBE), was added and samples were vortexed for 10 minutes, then centrifuged for 5 min (4 °C, 10,000 rpm) using a Microfuge 22R Centrifuge (Beckman Coulter, Brea, CA). The supernatant was transferred to a fresh tube. The lipid extraction was repeated once with the MTBE. The MTBE fractions were pooled, dried under argon gas, and stored at −80 °C.

### UHPLC-MS Untargeted Lipidomics

Extracts were reconstituted in 150 µL of acetonitrile/isopropanol (7:3, *v/v*), followed by 1 min of vortexing and 5 min centrifugation (4 °C, 14,000 rpm) using a Microfuge 22R Centrifuge (Beckman Coulter). Extract components were separated with a C8 column (Acquity Plus BEH, Waters, Milford, MA, 100 mm × 2.1 mm, 1.7 µm particle size) on Agilent 1290 UHPLC coupled to Agilent 6545 quadrupole time-of-flight mass spectrometer (Agilent Technologies, Santa Clara, CA). Samples were ionized using ESI probe operated in positive mode, followed by negative mode. Chromatographic separation conditions and source ionization parameters were shared previously [17]. The injection volume was 5 µL. Quality control samples (QC) were analyzed every 6th injection to correct for instrument performance. Blanks were analyzed before running the samples and analyzed every 25^th^ injection to remove persistent contaminant features from the data. Iterative MS-MS analysis on one QC was performed to capture MS2 data for assistance with metabolite identification.

### Lipidomics Data Processing

Raw UHPLC-MS data was processed using Agilent Profinder (version 10.0) to extract and align molecular features, and group by metabolite, using the following parameters: noise ≥ 10000 counts, retention time tolerance of 0.1 min, with a 10 ppm cutoff set for binning and alignment. Post-processing filtration was applied to eliminate metabolites with intensity< 30000 counts and to eliminate metabolites present in the process blanks. The Lipid Annotator software was used to process the iterative MS/MS files to generate a database, in order to annotate the previously extracted compounds using Agilent Mass Profiler Professional software.

### Lipid Peroxidation

Approximately 50 mg mammary gland (n=4-5/group) from day 28, 56, and 120 was added to 1X PBS with 0.5M butylhydroxytoluene and homogenized using Precellys Lysing Kit and Precellys Evolution Homogenizer. Samples were centrifuged and supernatant collected into new tubes. Malondialdehyde (MDA) protein adducts were measured using OxiSelect MDA Adduct Competitive ELISA kit (Cell Biolabs, Inc., San Diego, CA) per instructions. A four-parameter logistic regression curve analyzed results.

### Mitochondria and Whole Cell Reactive Oxygen Species

Mammary gland without lymph node from days 28, 56, and 120 postpartum (n=4-5/group) were digested with Type 1 Collagenase (Worthington Biochemical, Lakewood, NJ) and hyaluronidase (Millipore-Sigma) in Hank’s Balanced Salt Solution (HBSS) (Gibco, Evansville, IN) with 2% FBS. Cells were counted via Luna II (Logos Biosystems, Annandale, VA). The redox-sensitive fluorochrome 5-(and 6)-chloromethyl-2′,7′-dichlorodihydroflurescein diacetate dye (CM-H_2_DCFDA) (Invitrogen) was used to measure the intracellular reactive oxygen species. Mouse MG (n=5/group) cells were treated with Mitosox-Red and intracellular ROS concentration was evaluated at 6 and 24 h. In total, 5 × 10^4^ treated/untreated CD34+ cells were loaded with 2 μM CM-H_2_DCFDA for 20 min at 37 °C. Before analysis, the cells were removed from loading buffer and incubated in growth medium for 30 min at 37°C. Data acquisition and analysis were performed using a Cytek Northern Light (Becton Dickinson, BD, Franklin Lakes, NJ). At least 100,000 events were detected for each sample to guarantee statistical significance. Data was analyzed by FlowJo version 10.8 (Becton Dickinson).

### Seahorse Analysis

Mammary gland without lymph nodes from days 28, 56, and 120 postpartum (n=3-6/group) were digested with Type 1 Collagenase and hyaluronidase in HBSS with 2% FBS. Cells were counted via Luna II. A total of 50,000 cells from each sample were plated in triplicate on Seahorse cell culture plates and incubate at 37°C for four hours. Cells were analyzed by Seahorse Bioanalyzer XFe24 (Agilent Technologies) using XF Cell Mito Stress Test Kit (Agilent Technologies #103015-100). Protein was extracted and quantified using Pierce BCA assay as previously described. Results were normalized to protein quantity.

### Ingenuity Pathway Analysis of Pregnancy Associated Breast Cancer Samples

“Molecular Signature of Pregnancy Associated Breast Cancer (PABC)” dataset was identified on National Center for Biotechnology Information Gene Expression Omnibus (NCBI GEO #GSE31192). Gene expression profile of laser captured tumor epithelial cells and stroma were made publicly available by Harvell et. al [18]. Affymetrix Human Genome U133 Plus 2.0 Array data were downloaded and subjected to Ingenuity Pathway Analysis, as with mouse experiments (above). PABC estrogen receptor positive (ER+) and PABC estrogen receptor negative (ER-) tumor gene expression was evaluated for differential metabolic pathways via Ingenuity Pathway Analysis.

### Statistical Analysis

Data analyses were completed using statistical software SAS (SAS Institute Cary, North Carolina) and Prism (GraphPad, San Diego, CA). Data was analyzed at individual timepoints between AI and GI. Unless specified above, other comparisons were made using a two-sample t-test. The significance level for all analyses was set a priori at p<0.05, either for single comparisons or after Holm’s adjustment for multiple comparisons.

## 3. Results

### Day 28 AI vs GI Comparison of Mammary Glands

Previous findings suggested that while GI mice were still housed with offspring, the AI and GI mammary glands were histologically similar on day 28 postpartum [10,11]. Based on these findings, we first compared transcriptome of mammary glands harvested on day 28 via Ingenuity Pathway Analysis (**Figure 1b**). AI glands had significant upregulation of multiple metabolic pathways (p<0.05): oxidative phosphorylation, TCA cycle, mitochondrial protein import, mitochondrial fatty acid β-oxidation, gluconeogenesis, cristae formation, fatty acyl-CoA biosynthesis, glycolysis, mitochondrial translation, and acetyl-CoA biosynthesis. While not directional, lactose synthesis was significantly different between AI and GI glands. Validation was performed on genes that were differentially expressed with a greater than or equal to two fold change (**Supplementary Figure 2**) via qPCR between groups including (all p<0.05): *ACLY, ACSS, ELOVL3, GLUT5, CIDEA, Gpnmb2, ATP6v0d2, SLC25a1*, and *ACSM3*.

Mitochondrial complexes were evaluated via western blot to assess mitochondrial differences underlying these findings (**Figure 1c**). The quantification revealed AI glands had significantly higher concentration of complex I (p=0.0139) and complex IV (p=0.0083) than GI glands. AI glands had a trend for higher levels of complex II (p=0.075), complex III (p=0.1986), and complex V (p=0.1759) than GI glands. Untargeted lipidomics profiling indicated evidence of oxidative stress impacting several lipid species between groups with AI glands having an upregulation (**Figure 1d-h**) of oxidized polyunsaturated fatty acids commonly found in cell membranes including two sphingomyelin species (i.e., d18:2(4E,14Z)/18:1(9Z)-O(12,13); p=0.0030 and d18:1/18:1(9Z)-O(12,13); p=0.0064). AI glands had significantly higher concentrations of lipids with prostaglandin J2 (PGJ2) as part of the structure, including ceramide (t18:0/PGJ2; p=0.0141) and a diacylglycerol (PGJ2/18:0; p=0.0103). AI glands had an increase in phosphatidylcholine (18:1/18:2; p=0.0115), a common glycerophospholipid associated with pregnancy [19]. Interestingly, GI glands showed an upregulation of a ceramide phosphoethanolamine (d14:2(4E,6E)/16:0(2OH); p=0.001) and triglyceride (18:1/18:2/20:3; p=0.0205), commonly found in breastmilk (**Supplementary Figure 3c-e**) [19].

### Day 28 AI vs 56 GI

Extended weaning in mice has shown offspring will continue to breastfeed from the mother even though majority of their nutrition comes from solid food [15,16]. As suckling plays a major role in hormones that control milk protein synthesis [20], we examined gene expressions of two major milk proteins overtime in both AI and GI glands. Casein beta 2 (CSN2) decreased in AI glands the day after offspring removal (day 8.5) and continued to decreased overtime while GI glands had an increase in CSN2 expression on day 12 postpartum that began gradually decreasing on day 17 postpartum (**Figure 2a**). Comparison of CSN2 on day 28 showed that GI glands had a 10.5 fold increase in expression compared to AI glands (**Figure 2b**). Similar effects of AI and GI are shown with whey acidic protein (WAP). Overall gene expression of WAP on day 28 is lower in AI and GI glands compared CSN2 (**Figure 2c**). However, GI glands have a 47.5 fold higher expression of WAP than AI glands (**Figure 2d**). Based on the presence of milk lipid metabolites, expression of milk protein genes, and literature evidence of offspring breastfeeding past normal weaning time frames, we decided comparing glands on day 28 was not an accurate representation of metabolic changes induced by differences of involution. We examined metabolic differences between AI and GI glands based on time since removal of offspring that equated to day 28 for AI glands (21 days post weaning) and day 56 for GI glands (25 days post weaning).

**Figure 2.**
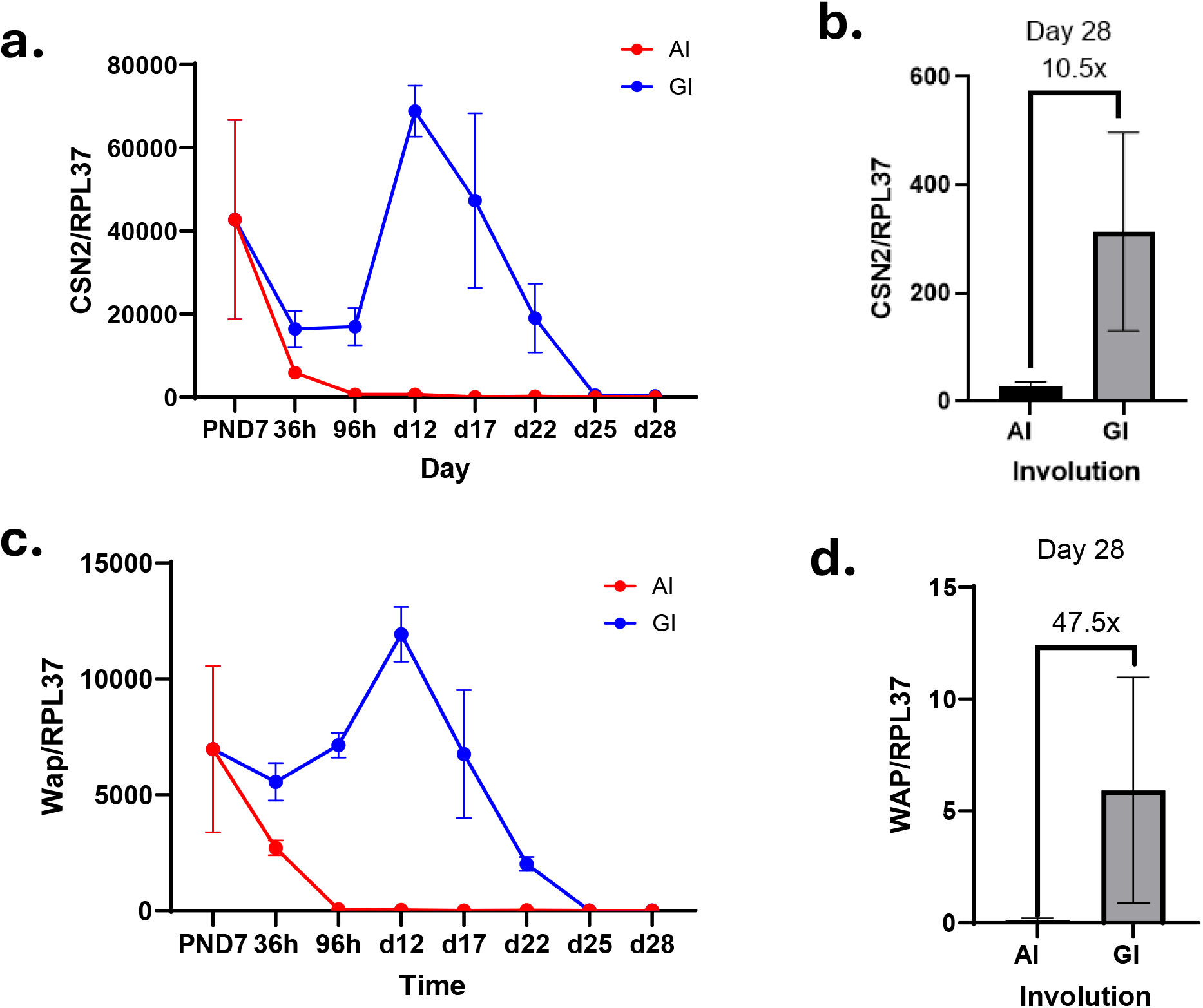
Analysis of Milk Protein Genes Overtime Between AI and GI Mammary Glands. **a**. CSN2 qPCR gene expression overtime **b**. CSN2 gene expression day 28. **c**. WAP qPCR gene expression overtime **d**. WAP gene expression day 28. AI= abrupt involution. GI=gradual involution. Data presented as mean ± SEM.

AI mammary glands upon Ingenuity Pathway Analysis (**Figure 3a**) had a significant upregulation (p<0.05) of numerous metabolic pathways compared to GI glands including glucose metabolism, insulin signaling, glycerophospholipid metabolism, cholesterol synthesis, mitochondria complex I and IV, mitochondria protein import, mitochondria biogenesis, and oxidative phosphorylation. On the contrary, AI mammary glands had a significant downregulation (p<0.05) compared to GI glands of complex III, glycolysis, and adipogenesis. While transcriptomic analyses indicated an increase in oxidative phosphorylation and mitochondrial complexes, there was no significant difference (**Figure 3b**) in complex I (p=0.4388), II (p=0.2648), IV (p=0.0724), or V (p=0.2633) between groups. GI had a significantly higher concentration of Complex III than AI (p=0.0345). To examine functional differences between AI and GI glands, Seahorse assay was used. Oxygen consumption rates were undetectable in AI glands unlike GI glands despite identical conditions. IPA indicates significant upregulation of several cellular stress pathways in AI glands that may impair functional analysis (**Figure 3a**): cellular response to hypoxia, cellular response to heat stress, cellular response to mitochondrial stress, NRF-2 mediated oxidative stress response, and detoxification of reactive oxygen species. When measured by flow cytometry, AI had significantly higher levels of mitochondrial oxidative stress (p=0.0002; **Figure 3c**).

**Figure 3.**
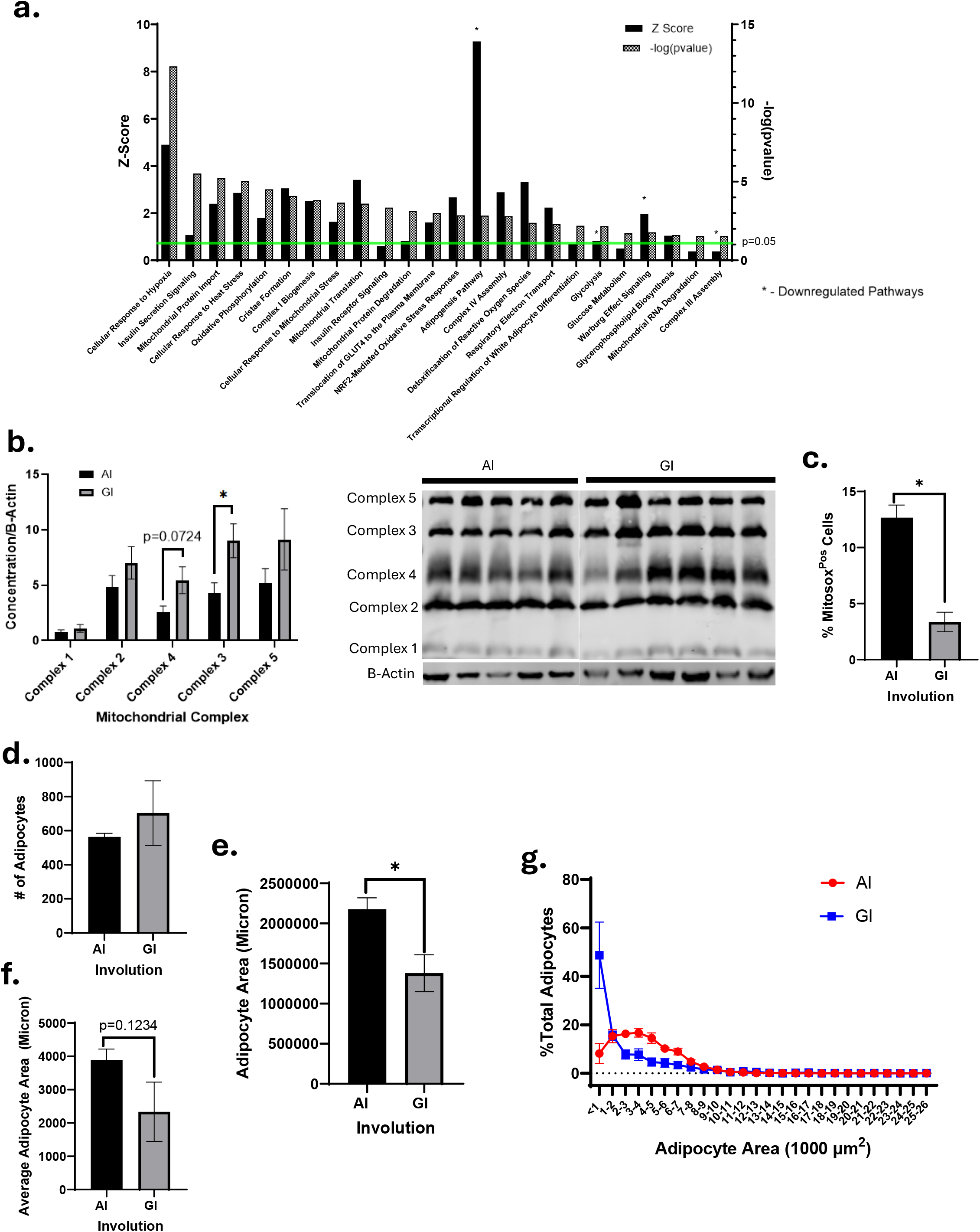
Metabolic Comparison of Day 28 AI and Day 56 GI Mammary Glands. **a**. IPA Analysis Pathways Impact by AI **b**. Western blot mitochondria complexes **c**. Mitosox flow cytometry analysis **d**. Average number of adipocytes **e**. Total adipocyte area in mammary glands **f**. Average adipocyte area **g**. Percent of adipocyte sizes by group. AI= abrupt involution. GI=gradual involution. Data presented as mean ± SEM. *depicts significant differences (p<0.05) between AI and GI glands.

To examine lipid metabolic pathways, adipocyte population of AI and GI glands were analyzed (**Figure 3d-g**). AI mammary glands had a significantly larger area of adipose tissue than GI glands (p=0.0261). AI and GI glands did not have significantly different number of adipocytes (p=0.4226), however AI mammary glands had a trend for larger average size of adipocytes than GI glands (p=0.1234). Adipocyte populations were evaluated by size per group with the GI group having more adipocytes <1000 microns and AI having more adipocytes greater than 1000 microns. These results suggest that adipocytes in AI glands may undergo hypertrophy in repopulation compared to GI glands.

### AI Causes Long-term Metabolic Dysregulation

Given substantial differences in mammary gland metabolism and mitochondrial oxidative stress, AI and GI glands were evaluated at day 120 (long-term effect). IPA of day 120 transcriptomic data revealed several metabolic pathways impacted by AI compared to GI (**Figure 4a**). AI glands showed significant downregulation of metabolic pathways (p<0.01 for all) including oxidative phosphorylation, mitochondrial translation, glycolysis, gluconeogenesis, mitochondrial protein import, serine and glycine biosynthesis, cristae formation, and mitochondrial biogenesis. There was upregulation of mitochondrial dysfunction pathway. There were no significant differences of mitochondrial complexes (p>0.05) (**Figure 4b**).

**Figure 4.**
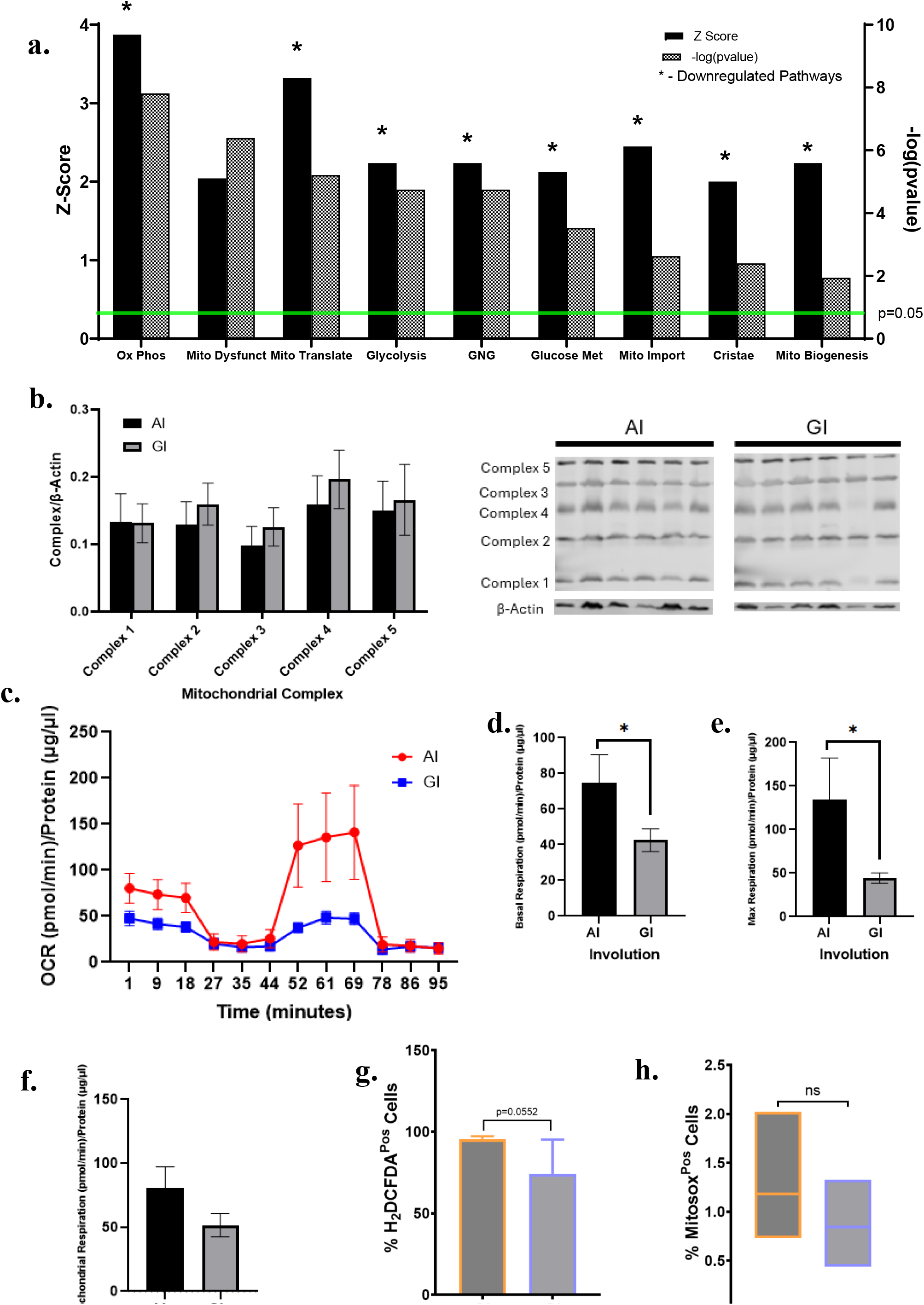
Day 120 Post-Partum Enrichment of Mitochondrial Dysfunction in AI Glands. **a**. Ingenuity Pathway Analysis of significantly impacted metabolic pathways in AI glands. **b**. Western blot of mitochondrial complexes. **c**. Seahorse functional analysis of AI and GI mammary gland cells. **d**. Basal respiration of mammary gland cells. **e**. Maximum respiration potential of mammary gland cells. **f**. Mitochondrial respiration of mammary gland cells. **g**. Flow cytometry of whole cell oxidative stress (H2DCDFA). **h**. Flow cytometry of mitochondria oxidative stress (MitoSox). AI= abrupt involution. GI=gradual involution. Data presented as mean ± SEM. *depicts significant differences (p<0.05) between AI and GI glands.

Functional analysis showed AI glands had significantly higher basal respiration (p=0.0011) and maximal respiration (p=0.0027) compared to GI glands (**Figure 4c-e**). There was no significant difference between the two groups for mitochondrial respiration (p=0.2190; **Figure 4f**). Analysis of H2DCFDA and MitoSox revealed no significant differences between the groups, although AI glands had a trend for higher MitoSox levels than GI (**Figure 4g-h**). Lipidomics analysis showed the relative abundance of oxidized phosphatidic acid (30:3;O2) was significantly higher in AI glands than GI (p=0.0476; **Figure 4**). While oxidation of metabolite may suggest oxidative stress, phosphatidic acid has been implicated in metastatic ability of breast cancer cells [21]. Collectively, day 120 AI metabolic profiling demonstrated that transcriptional profiles were largely opposite of day 28, indicating potential mitochondrial dysfunction in the AI glands after 4 months.

### Metabolic Pathways in Human ER-Negative Pregnancy-Associated Breast Cancer Similar to Day 120 AI Mammary Glands

To translate our findings and understand metabolic differences in breast tumors following pregnancy in humans, a dataset examining gene expression of tumors categorized as pregnancy associated breast cancer (PABC) was analyzed [18]. Epithelial and stroma tumor gene expressions were compared and analyzed by Ingenuity Pathway Analysis software to identify differentially enriched metabolic pathways (**Figure 5a**). PABC ER-tumors displayed a significant enrichment (all p<0.05) in several metabolic pathways such as mitochondrial dysfunction, triglyceride metabolism, mitochondrial biogenesis, and cholesterol biosynthesis. PABC ER-tumors had significantly reduced pyruvate metabolism, fatty acyl-CoA biosynthesis, signaling by insulin receptor, mitochondrial translation, and integration of energy metabolism. There were significant differences between PABC ER- and ER+ tumors with respect to insulin secretion signaling, amyloid processing, serine biosynthesis, and mitochondrial division signaling. Several pathways altered in PABC ER-tumors overlap with pathways in day 120 AI glands. These results were compared to pathways impacted by AI on day 120 (**Figure 5b**). Several metabolic pathways were found down regulated in both PABC ER-tumors and AI MG including mitochondrial translation, oxidative phosphorylation, mitochondrial protein import, and cristae formation. Mitochondrial dysfunction was upregulated in both PABC ER- and AI MG.

**Figure 5.**
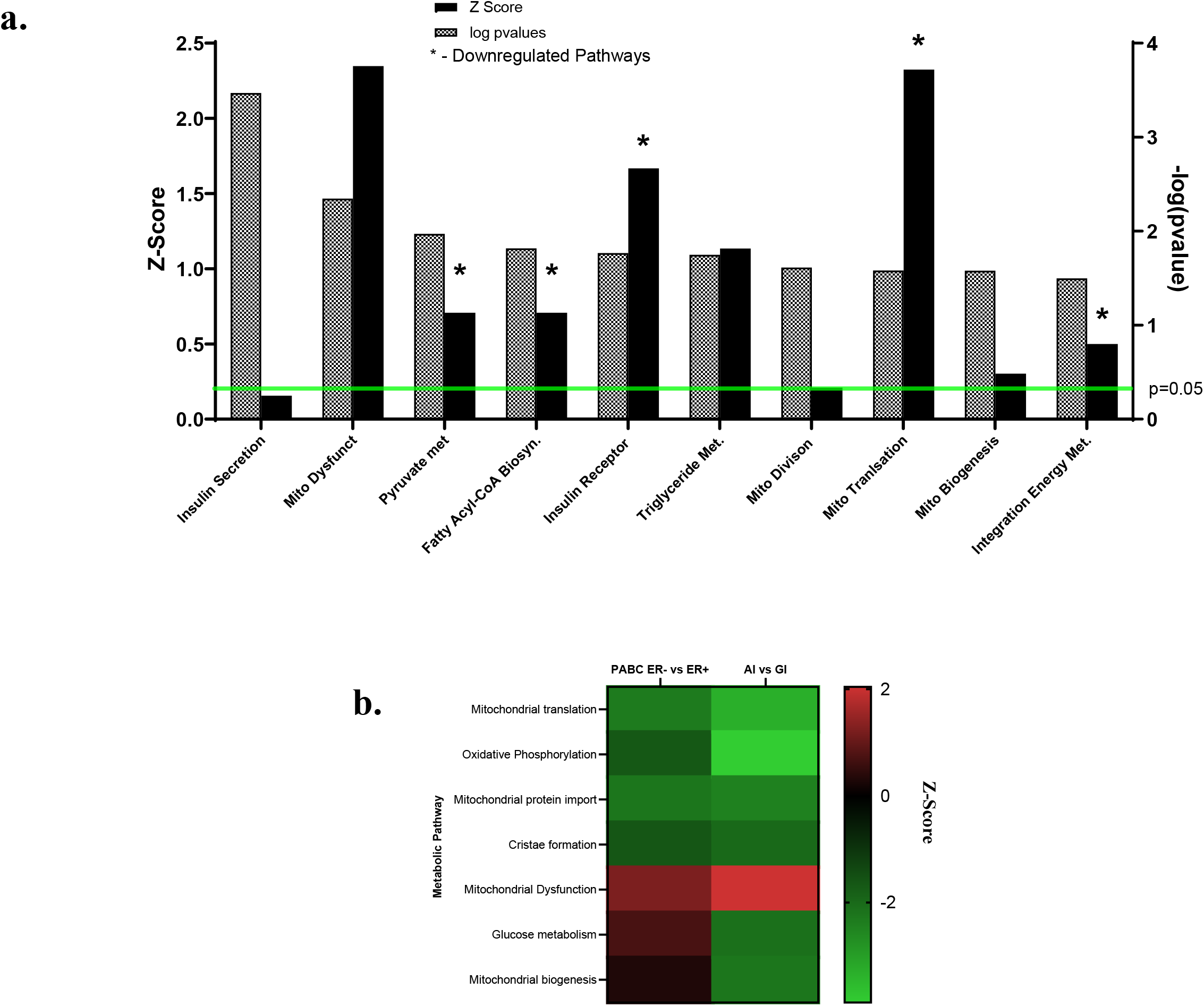
Ingenuity Pathway Analysis of ER- and ER+ Pregnancy Associated Breast Cancer samples. **a**. Significantly altered metabolic pathways in PABC ER-tumors compared to PABC ER+ tumors. **b**. Comparison of metabolic pathways between human ER-PABC and day 120 AI mouse mammary glands.

### Blocking of Estrogen Receptor with Tamoxifen Study

Literature on involution has shown that estrogen return during AI leads to neutrophil infiltration and extracellular matrix remodeling that directly and indirectly promotes adipocyte re-differentiation as previously discussed [12]. In addition to metabolic pathways impacted by AI shown in **Figure 3a**, IPA day 28 AI glands (**Supplementary Figure 3a**) showed upregulation of estrogen signaling, neutrophil degranulation, and several RNA/DNA synthesis pathways compared to GI glands. We previously reported H-scores for ERα in AI and GI mammary glands at each respective timepoint (day 28 and 56) [10]. Comparison of day 28 AI glands and day 56 GI glands showed AI glands had significantly greater ERα positivity via IHC in epithelial populations and stromal compartments than day 56 GI glands (p<0.0001 for both) (**Supplementary Figure 3b**). As previously mentioned, AI glands had greater adipocyte area and trend for larger adipocytes than GI glands. Based on the literature showing estrogen’s impact on adipocyte repopulation and upregulation of estrogen mediated signaling and neutrophil degranulation, we hypothesized that the return of estrogen during AI leads to the metabolic changes that could support altered RNA metabolism and cell proliferation.

To test this hypothesis, a subset of FVB/n mice underwent breeding and AI as previously described (**Figure 1a**). On day 8 postpartum, sustained release tamoxifen citrate (5mg) or placebo pellet were placed in the subscapular region and mice were harvested on day 28 postpartum. IPA showed tamoxifen (**Figure 6a**) treated mice had significant (p<0.05) down regulation of extracellular matrix organization, neutrophil degranulation pathways and upregulation of adipogenesis. Additionally, tamoxifen treated mice had upregulation (p<0.05) of oxidative phosphorylation, respiratory electron transport, complex I biogenesis, glucose metabolism, glycolysis, and downregulation of glycerophospholipid biosynthesis (**Figure 6a**). Western blot of mitochondrial complexes (**Figure 6b**) showed tamoxifen treated glands had significantly more complex I (p=0.0254) than placebo treated glands. While not significant, there was a trend for tamoxifen treated glands to have higher concentrations of complex IV (p=0.1780) and V (p=0.1022). There were no significant differences between groups for number of adipocytes, total adipocyte area, or average adipocyte size (**Figure 6 c-f**).

**Figure 6.**
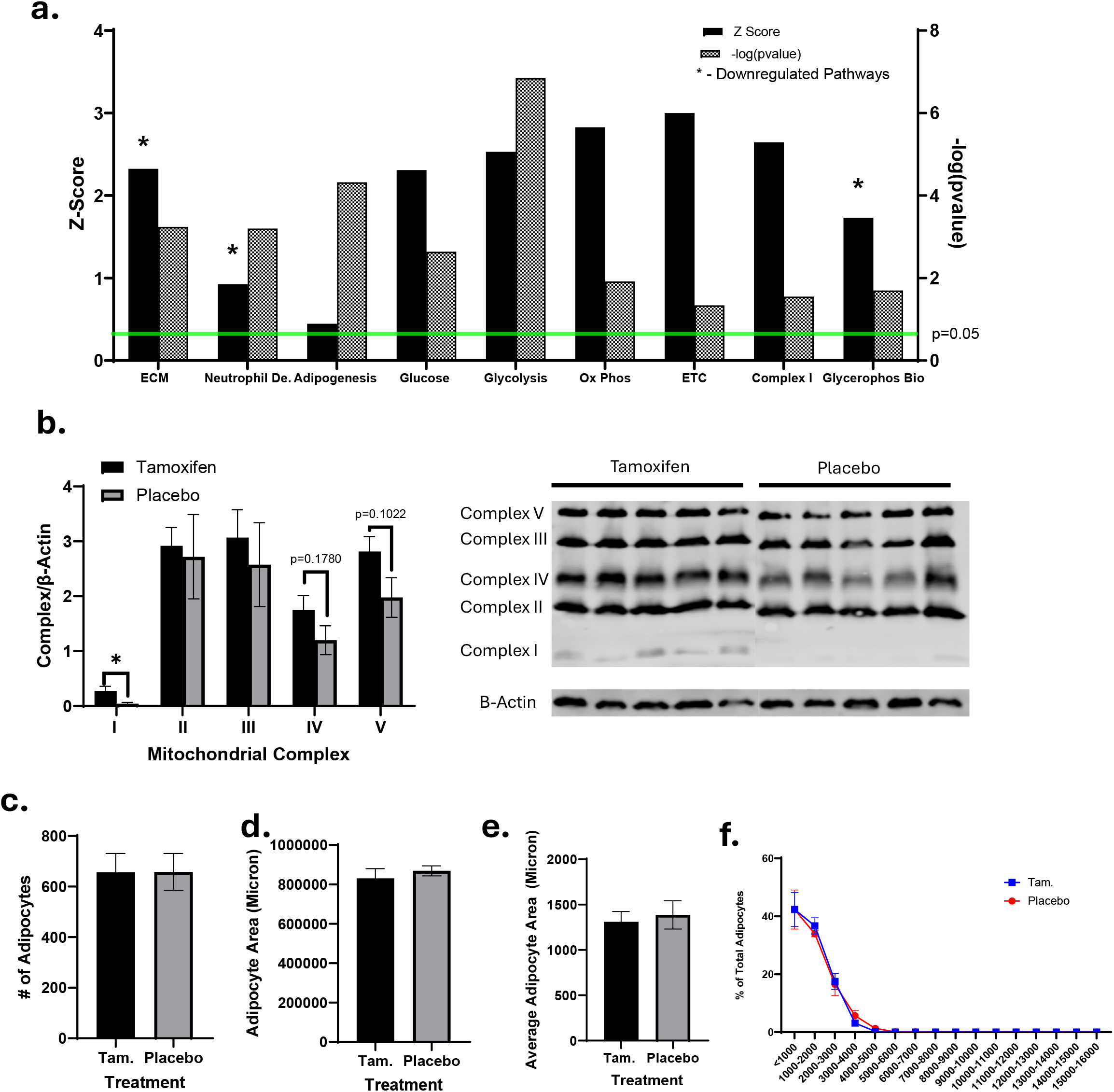
Metabolic Comparison of Day 28 Tamoxifen versus Placebo Mammary Glands. **a**. IPA Analysis of pathways impacted by tamoxifen compared to placebo **b**. Western blot mitochondria complexes **c**. Average number of adipocytes **d**. Total adipocyte area **e**. Average adipocyte area **f**. Percent of total adipocytes by size. AI= abrupt involution. GI=gradual involution. Data presented as mean ± SEM. *depicts significant differences (p<0.05) between AI and GI glands.

## 4. Discussion

To our knowledge, this is the first study to examine metabolic changes within the mammary gland of mice undergoing AI and GI. Expression of milk protein genes and presence of milk lipids during early analyses showcased that while AI and GI glands were histologically similar, the glands were not metabolically comparable. Comparison of day 28 AI and day 56 GI glands showed abrupt cessation of lactation led to upregulation of estrogen signaling, metabolic pathways associated with anabolism, and greater adipocyte area, along with elevated markers of mitochondrial oxidative stress and oxidized fatty acids. Tamoxifen treatment in AI mice resulted in metabolic outcomes and adipocyte repopulation similar to GI glands suggesting a role of estrogen in metabolic phenotypes observed in AI glands. Long-term transcriptomic analysis revealed downregulation of key metabolic pathways accompanied by signs of mitochondrial dysfunction in AI glands. Several metabolic pathways altered in day 120 AI glands were similar to PABC ER-tumors, suggesting a link of AI to TNBC through metabolism.

Lim et al. found during involution estradiol binds to estrogen receptor alpha (ERα) causing an increase in neutrophils and extracellular remodeling which indirectly cause adipocyte re-population during the second phase of involution (~day 14 postpartum). Adipocytes in AI glands expand after re-differentiation due to hypertrophy of milk lipids. Interestingly, our results two weeks later in day 28 AI glands show similar effects with upregulation of estrogen receptor signaling, neutrophil pathways, and larger adipocytes/adipocyte area. The AI glands have an increase in glucose metabolism with increased glucose uptake into adipocytes via GLUT4 though glycolysis is downregulated. Coupled with upregulation of RNA and DNA synthesis and mitochondrial biogenesis pathways, this suggests the divergent of metabolism towards cell growth and proliferation. Research has shown that rapid adipocyte re-population and hypertrophy cause increased adipocyte glucose uptake, mitochondrial biogenesis, oxidative phosphorylation, protect against oxidative stress, and promote cell proliferation similar to previously shown outcomes [22–24].

GI glands had a downregulation of estrogen signaling, upregulation of adipogenesis, and a subsequent different metabolic phenotype compared to AI. Results of anti-estrogen therapy, tamoxifen, in AI glands mimicked several findings in GI glands including upregulation of glycolysis and adipogenesis, and down regulation of neutrophil pathways. Tamoxifen treatment did not replicate the entirety of findings in GI glands. As tamoxifen treated AI glands show upregulation of adipogenesis compared to adipocyte hypertrophy, the fate of milk lipids present at the time of weaning is not clear. This may contribute to metabolic differences between tamoxifen treated AI and GI glands. With key differences between GI (and AI tamoxifen treated) and AI glands, it could be inferred that the resurgence of estrogen imparted in the glands cause differences in adipocyte repopulation; a critical point in determining the metabolic phenotype and cell proliferation.

Long-term timepoint reverses transcriptional changes originally shown on day 28 AI glands. There is downregulation of energy pathways and upregulation of mitochondrial dysfunction. While mitochondrial complexes are not different, all mitochondrial complexes are lower in AI than GI. PABC ER-tumors showed similar affected metabolic pathways to AI glands. Both AI glands and PABC ER-tumors displayed upregulation of mitochondrial dysfunction and downregulation of oxidative phosphorylation. PABC tumors had higher amounts of cell death and extracellular matrix remodeling similar to previous findings of AI. While not evaluated in PABC tumors, AI glands showed similar elevation of phosphatidic acid as metastatic MDA-MB-231 TNBC cell lines [25]. These results provide potential links between AI and ER-breast cancer, that have been linked in epidemiological studies.

This study has acknowledged strengths and limitations. This research focused on metabolic changes overtime related to AI and GI, but earlier timepoints need to be evaluated to understand the short-term impact that causes downstream, long-term effects. Potential mechanisms of estrogen signaling on adipocyte repopulation provided critical information regarding estrogen’s role in mammary gland biology, metabolic outcomes, and subsequent long-term impact of its effects. Further studies need to be conducted to examine the differences in estrogen signaling between AI and GI glands and how the adipocyte repopulation process leads to metabolic differences.

## 5. Conclusion

This research study shows tissue specific metabolic effects of AI over time and the potential role of estrogen signaling. Additionally, AI leads to long-term mitochondrial dysfunction. These alterations closely mirror metabolic signatures observed in PABC ER-tumors. Together, these findings support a potential link between lack of breastfeeding and increased breast cancer risk.

## 7. Contributions

KO-Conceptualization, Data Curation, Formal Analysis, Investigation, Project Administration, Resources, Supervision, Visualization, Funding, Writing. NS – Conceptualization, Investigation, Project Administration, Writing. GS – Investigation. AZ – Investigation. MB – Investigation, Data Curation, Writing. RKS – Investigation, Validation. SA – Investigation. AG – Investigation. DB – Validation, Curation, Investigation. JW – Investigation, Writing. XZ – Formal Analysis. RK – Methodology, Validation, Curation, Investigation, Writing. ES – Supervision, Writing. KIS – Methodology, Resources, Validation, Supervision, Writing. RG – Funding, Writing. SM – Conceptualization, Project Administration, Supervision, Resources, Funding. BR – Conceptualization, Project Administration, Supervision, Funding. DS – Project Administration, Supervision, Funding, Writing.

## 8. Acknowledgements

We thank Genomics Shared Resource at The Ohio State University Comprehensive Cancer Center (OSU CCC), Columbus, OH for conducting the Affymetrix Clairom D analyses. This work was supported in part by the OSUCCC and the National Institutes of Health grant P30 CA16058. The content is solely the responsibility of the authors and does not necessarily represent the official views of the National Institutes of Health.

We would like to acknowledge Dr. Bhuvaneswari Ramaswamy who passed away from breast cancer in July 2024 during this research. Dr. Ramaswamy spearheaded this work in involution, and we want to carry on her legacy.

## 9. Data Statement

Data available upon request.

## 11. Ethics Statement

All experiments were conducted according to National Institute of Health Guide for the Care and Use of Laboratory Animals. All protocols were approved by the Ohio State Institutional Animal Care and Use Committee

