## Supplementary Figures for "Estrogen Signaling During Abrupt Involution Leads to Long-Term Metabolic Dysfunction Similar to Estrogen Receptor Negative Breast Cancer"

Supplementary Table 1. qPCR Primer Sequences

| Gene Name | Forward Sequence | Reverse Sequence |
|---|---|---|
| ACLY | GTCCCAAGTCCAAGATCCCTG | TGTGATCCCCAGTGAAAGGG |
| ELOVL3 | CATGAATTTCTCACGCGGGTT | TTGCTTGAGGCCCACTGTAA |
| Acss2 | AGGTGACCAAGTTCTACACGGC | GTTGATGGGTTCACCTACTGTGC |
| GLUT5 | ACAGCTGGCACTTTGAGGAG | TGCTGCATGAACTCTGAGGG |
| SLC25a1 | CCCCTTCATTCAAATGGGCCT | AATGTCCTGCCCTTGGTCTC |
| CIDEA | CATACATCCAGCTCGCCCTT | TACTACCCGGTGTCCATTTCTG |
| Gpnmb | CCCCTTCGCCTTCGACTC | CTTCCAGGATCCCCTCTACA |
| ATP6v0d2 | GGGCCAGTGTTCAGTTGCTA | AGTCCGTGGTCTGGAGATGA |
| CPT2 | CAACTCGTATACCCAAACCCAGT | GTTCCCATCTTGATCGAGGACAT |
| Acsm3 | CCGGATGCTTGTTCAGAATGAC | TCCACAGATCAGCACCGTTT |
| GLUT4 | TATTGCAGCGCCTGAGTCTT | TTCAATCACCTTCTGTGGGGC |
| PGC1α | AAGGTCCCCAGGCAGTAGAT | CATAGCTGTCGTACCTGGGC |
| Chrebp | CGACACTCACCCACCTCTTC | TTGTTCAGCCGGATCTTGTC |
| APP | GACCACTCGACCAGGTTCTG | GATGATGGCGCCTTTGTTCG |
| Adam10 | TGCCAAACGAGCAGTCTCAC | TCGTAGGTTGAACTGTCTTCCA |
| TIMM23 | GGTTAGCTGGCTTCTTCGGA | TTCATACCAGTCAGCGGGAC |
| TOMM40 | AGCAACCGTTTCCAGGTGAC | TTGTCCATGTCACCCACCAG |
| RPL37 | AGAAGCAAGATGACGAAGGGAA | CTTGGGTTTCGGCGTTGTTC |

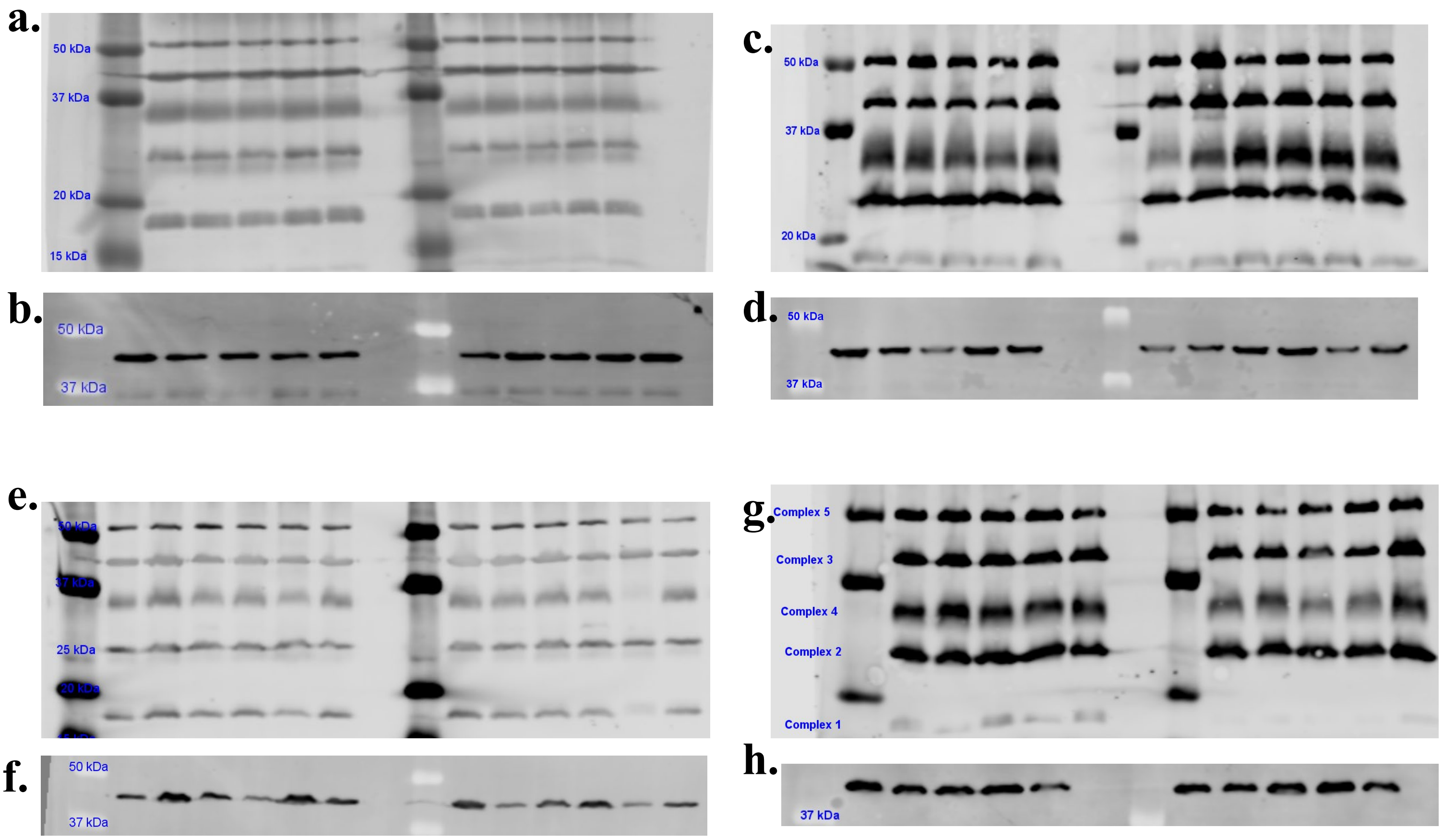


**Supplementary Figure 1.** *Full Western Blots Analyzed.* **a**. Mitochondrial Complexes day 28 AI vs GI. b. β-Actin day 28 AI vs GI. c. Mitochondrial Complexes day 28 AI vs day 56 GI. d. β-Actin day 28 AI vs day 56 GI. d. Mitochondrial complexes day 120 AI vs GI. e. β-Actin day 120 AI vs GI. f. Mitochondrial complexes tamoxifen vs placebo day 28. g. β-Actin tamoxifen vs placebo day 28.

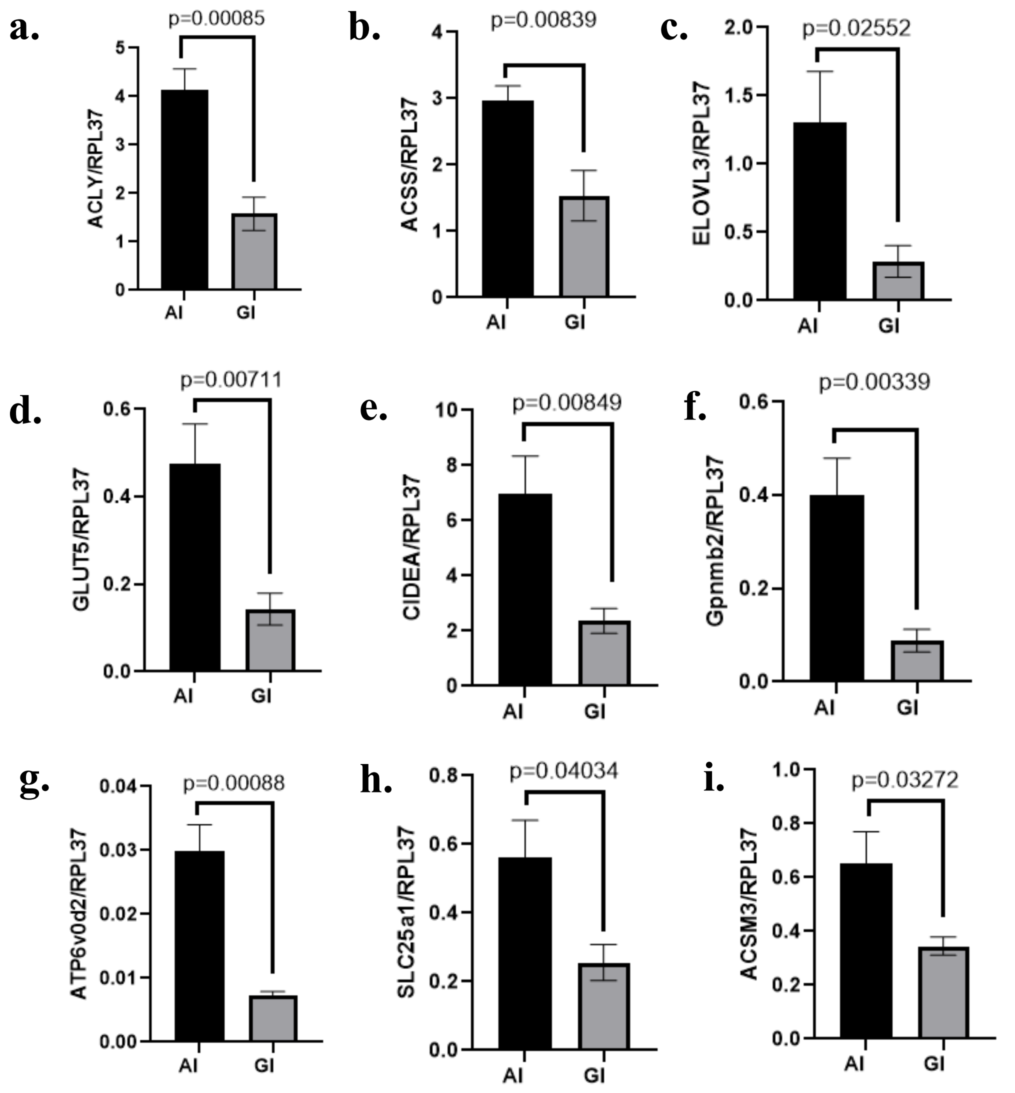


**Supplementary Figure 2.** *Day 28 Upregulated Metabolic Genes in AI Mammary Glands Validated via qPCR.* **a**. ACLY **b**. ACSS **c**. ELOVL3 **d**. GLUT5 **e**. CIDEA **f**. Gpnmb2 **g.** ATP6v0d2 **h.** SLC25a1 **i.** ACSM3. Genes were normalized to housekeeping gene RPL37. AI= abrupt involution. GI=gradual involution. Data presented as mean ± SEM. $^*$depicts significant differences ($p<0.05$) between AI and GI glands.

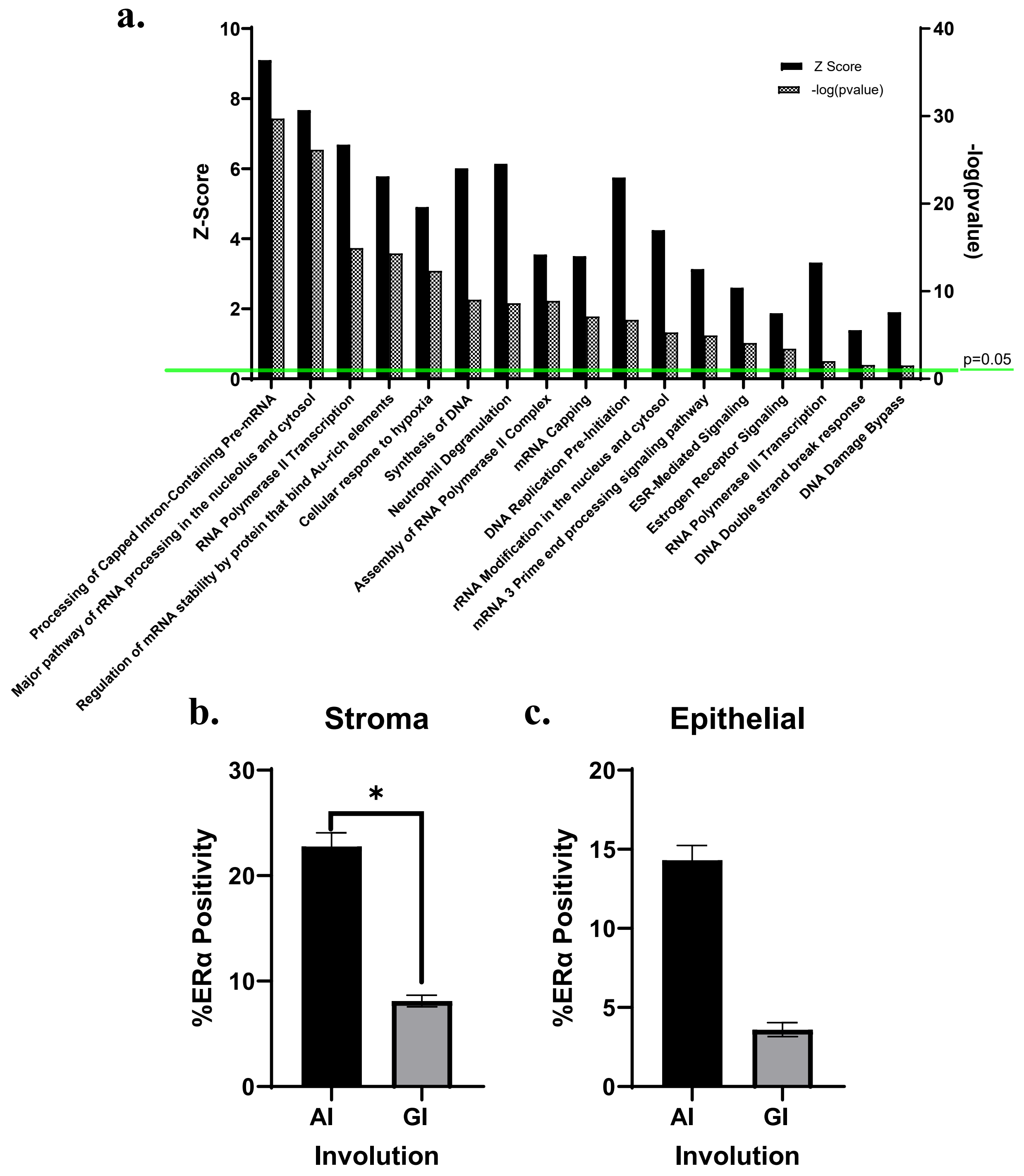


**Supplementary Figure 3.** *Estrogen Signaling in Day 28 AI vs Day 56 GI Mammary Glands* **a**. Upregulated pathways in AI glands from IPA Analysis **b**. ERα positivity in stroma via IHC **c**. ERα positivity in epithelial cells via IHC. AI= abrupt involution. GI=gradual involution. Data presented as mean ± SEM. *depicts significant differences ($p<0.05$) between AI and GI glands.
